# A lentiviral fluorescent reporter to study circadian rhythms in single cells

**DOI:** 10.1101/2025.06.30.661926

**Authors:** Christian H. Gabriel, Luis Lehmann, Joana Ahlburg, Achim Kramer

## Abstract

Circadian rhythms—self-sustained, ∼24-hour oscillations in transcript and protein levels—are generated by a cell-autonomous molecular clock. These rhythms shape how individual cells respond to external signals, influencing key decisions such as differentiation and apoptosis. However, current tools for visualizing circadian rhythms at the single-cell level often rely on genomic engineering and clonal expansion, limiting their accessibility and applicability. We present fluorescent circadian reporters based on the murine *REVERBα/Nr1d1* gene, delivered via lentiviral transduction and compatible with time-lapse single cell microscopy. These reporters produce oscillatory signals that depend on a functional circadian clock and can be used to determine a cell’s circadian dynamics parameters, such as phase and period. Their simple and efficient delivery makes them suitable for a wide variety of cell types, greatly expanding opportunities to study single-cell circadian dynamics and their impact across diverse biological processes and systems.

## Introduction

The environment of most organisms is shaped by the recurring changes in sunlight intensity between day and night. In response, the majority of species have evolved timekeeping mechanisms—circadian clocks—that drive rhythmic physiology, metabolism and behavior, enabling them to anticipate and adapt to these cycles. These circadian rhythms are self-sustained and persist in the absence of external cues. Circadian oscillations are evident at the organismal (e.g., sleep-wake cycles, hormone secretion (de Assis and Kramer 2024), immune activity (Scheiermann et al. 2013) and tissue level (e.g., rhythmic hepatic enzyme expression (Bolshette et al. 2023)). However, these rhythms originate from a molecular oscillator present in nearly every cell. Tissue- and organ-level rhythms emerge through paracrine and endocrine communication among these individual cellular clocks (Abraham et al. 2010; Finger et al. 2021). At the core of this oscillator is a transcriptional-translational feedback loop (TTFL). The BMAL1/CLOCK transcription factor dimer activates expression of core clock and clock-controlled genes, including those encoding PERIOD (PER1–3) and CRYPTOCHROME (CRY1–2) proteins. PERs and CRYs inhibit BMAL1/CLOCK, thereby repressing their own transcription until they degrade, allowing the cycle to restart (Finger and Kramer 2021). Another clock-controlled factor, REVERBα (encoded by *Nr1d1*), feeds back into the loop by rhythmically repressing *Bmal1* transcription (Preitner et al. 2002).

Studying circadian rhythms at the single-cell level—where they originate—offers deeper insights into the circadian network hierarchy and enables detection of phase-dependent variations in cellular responses, such as time-of-day sensitivity to drugs, toxins, or chemotherapeutics (Mihelakis et al. 2022; Ector et al. 2024). While many studies have relied on synchronized cell populations, relating the response of each single cell to its own internal clock allows for higher temporal precision with less experimental complexity, as cells in various phases can be analyzed within the same sample (Manella et al. 2021; Gabriel et al. 2024). However, tools for such single-cell circadian studies remain limited. Most approaches require longitudinal measurement of a reporter signal in individual cells, typically achieved through time-lapse microscopy using fluorescent circadian reporters. A common method involves genetically engineering cells or animals to express oscillating proteins like PERs or CRYs as fluorescent fusions (Gabriel et al. 2021; Smyllie et al. 2025), but this requires intensive genetic manipulation and clonal expansion. An alternative strategy employs a reporter plasmid driven by the rhythmically expressed *Nr1d1* locus, leading to oscillatory REVERBα-based fluorescence in transfected cells (Nagoshi et al. 2004). Despite offering insights into how the clock governs cell growth and differentiation (Zhang et al. 2022; Gutu et al. 2025) the use of this approach is constrained by the large plasmid size (∼18 kb), which reduces transfection efficiency and necessitates clonal selection. Consequently, only a few reporter cell lines based on this construct have been established over the past two decades.

To overcome these limitations, we generated fluorescent reporters of circadian rhythms that can be delivered by lentiviral transduction. They are based on the REVERBα reporter and allow the generation of reporter cells within a few days. The reporter signal oscillates in a circadian clock dependent manner and can be used to determine circadian dynamics parameters such as circadian phase and period at the single-cell level.

## Results

### Construction of a lentiviral fluorescent circadian reporter

To record circadian rhythms from single cells without the need for clonal isolation, we designed a lentiviral circadian fluorescence reporter plasmid based on the well-established REVERBα-Venus reporter (hereafter referred to as the ‘full-length reporter’). This reporter was generated by Nagoshi and colleagues two decades ago (Nagoshi et al. 2004). The reporter construct consists of the entire *Nr1d1*/REVERBα gene locus, spanning the region from approximately 6 kb upstream of the transcription start site (TSS) to 3.6 kb downstream of the last of eight exons, with a destabilized nuclear Venus protein (Venus-NLS-PEST) expression cassette replacing a region of exon 2-5 (Fig. 1A, yellow box), giving the construct a total size of approximately 18 kb.

**Figure 1:**
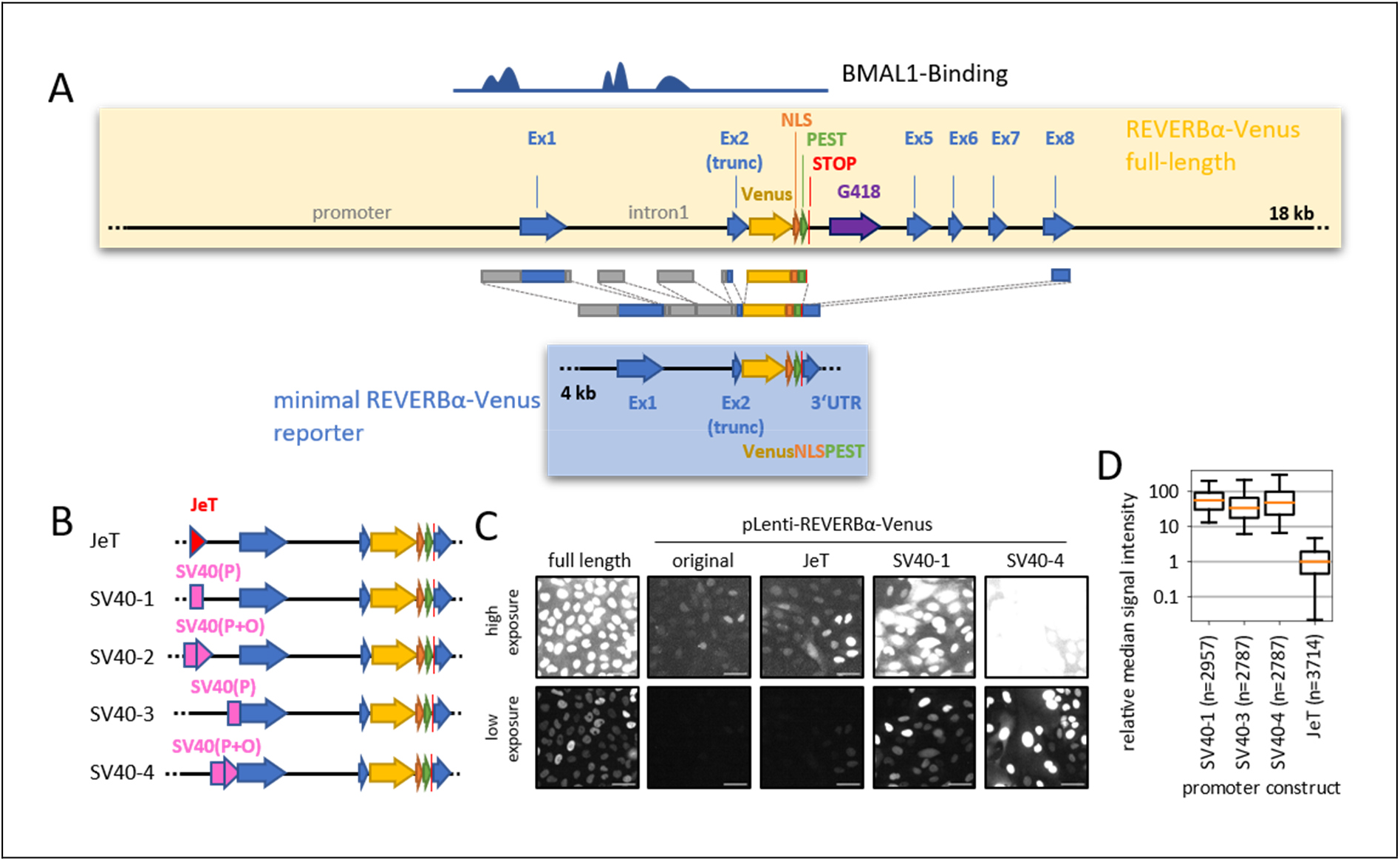
Construction of lentiviral fluorescent REVERBα reporters. **(A)** Schematic representation of the minimal REVERBα-Venus reporter (blue) and the full-length REVERVα-Venus construct (yellow) as previously described ((Nagoshi et al. 2004). BMAL1 binding regions are indicated based on data from (Koike et al. 2012). **(B)** Brighter reporter variants were generated by introducing either the synthetic JeT promoter, the SV40 minimal early promoter (P) or a combination of promoter and SV40 origin (P+O), positioned either proximal or distal to the REVERBα promoter. **(C)** Representative Venus fluorescence in U-2 OS cells either stably transfected with full-length REVERBα-Venus construct, or transduced with the indicated lentiviral reporter variants. Scale bar: 50 µm. **(D)** Background-subtracted median reporter fluorescence intensities per single-cell time series (68 h), normalized to median of JeT-construct. n = number of cells. Data representative of two experiments, whiskers = 5-95 %.

To fit into a lentiviral plasmid, we reduced the regulatory part of the reporter to the regions reported to be bound by BMAL1 (Chip-seq data from (Koike et al. 2012)), i.e., 600 bp of the proximal promoter and two regions within the first exon. Furthermore, we retained the sequence of the 3’-untranslated region (UTR), as these regions often contain sequences that regulate expression. Together with the Venus-NLS-PEST, this produced a reporter of ∼4 kb that fits a lentiviral backbone (Fig. 1A, blue box). The complete reporter sequence was synthesized and sub-cloned into a lentiviral expression plasmid containing a puromycin resistance cassette. The resulting ∼11 kb plasmid is within the upper size range usually regarded suitable for effective lentiviral packaging (Park 2007).

When we transduced this first lentiviral reporter into U-2 OS cells (an osteosarcoma and circadian model cell line), we observed fluorescence signals in the nuclei of the cells. This indicates that the Venus-NLS-PEST sequence is produced from the truncated reporter. However, the Venus signals were very close to the background level and approximately 50-100 times lower than those observed in a U-2 OS clone that stably expresses the full-length reporter (Fig. 1C). This suggests that the lentiviral reporter lacks regions from the *Nr1d1*-locus that, although not bound by BMAL1, play a role in regulating REVERBα expression levels.

We wondered whether we could enhance the expression of the reporter in order to facilitate detection and to enhance robustness against technical noise. To this end, we added either the synthetic JeT promoter (Tornoe et al. 2002), or parts of the SV40 early promoter (the SV40 minimal early promoter alone or in combination with the SV40 ori) either proximal or distal to the *Nr1d1* promoter region (Fig. 1B). The addition of the JeT promoter resulted in a 10-fold increase in signal (Fig. 1C), while the addition of SV40 promoter parts to an additional 30 – 50-fold increase (Fig. 1C-D). Interestingly, the fluorescence intensity of individual cells fluctuated substantially over the course of days (Supplementary Video SV1), suggesting that reporter expression is still subject to circadian regulation despite the increase caused by the constitutive promoters.

### Reporter signals are shaped by the circadian clock

Next, we aimed to establish whether these fluctuations are in fact caused by circadian regulation, i.e. whether they depend on an intact circadian clock. To this end, we introduced the lentiviral reporters into wild-type (WT) U-2 OS cells as well as cells in which the circadian oscillator is disrupted by knocking out both CHRYPTOCHROME proteins CRY1 and CRY2 (Bording et al. 2019). When we sorted the reporter signal from hundreds of single WT cells by their peak, oscillatory patterns emerged (Fig. 2A). These patterns aligned when the cells were synchronized with dexamethasone prior to recording (Fig. 2B), resulting in an oscillatory population signal (Fig. 2C). In contrast, oscillatory patterns were absent at both the single cell and population level when the same reporters were introduced into CRY-deficient cells (Fig. 2D-F). This demonstrates that the observed signal oscillations primarily depend on a functional circadian clock, suggesting that the reporters’ signals are driven by the core circadian mechanism.

**Figure 2:**
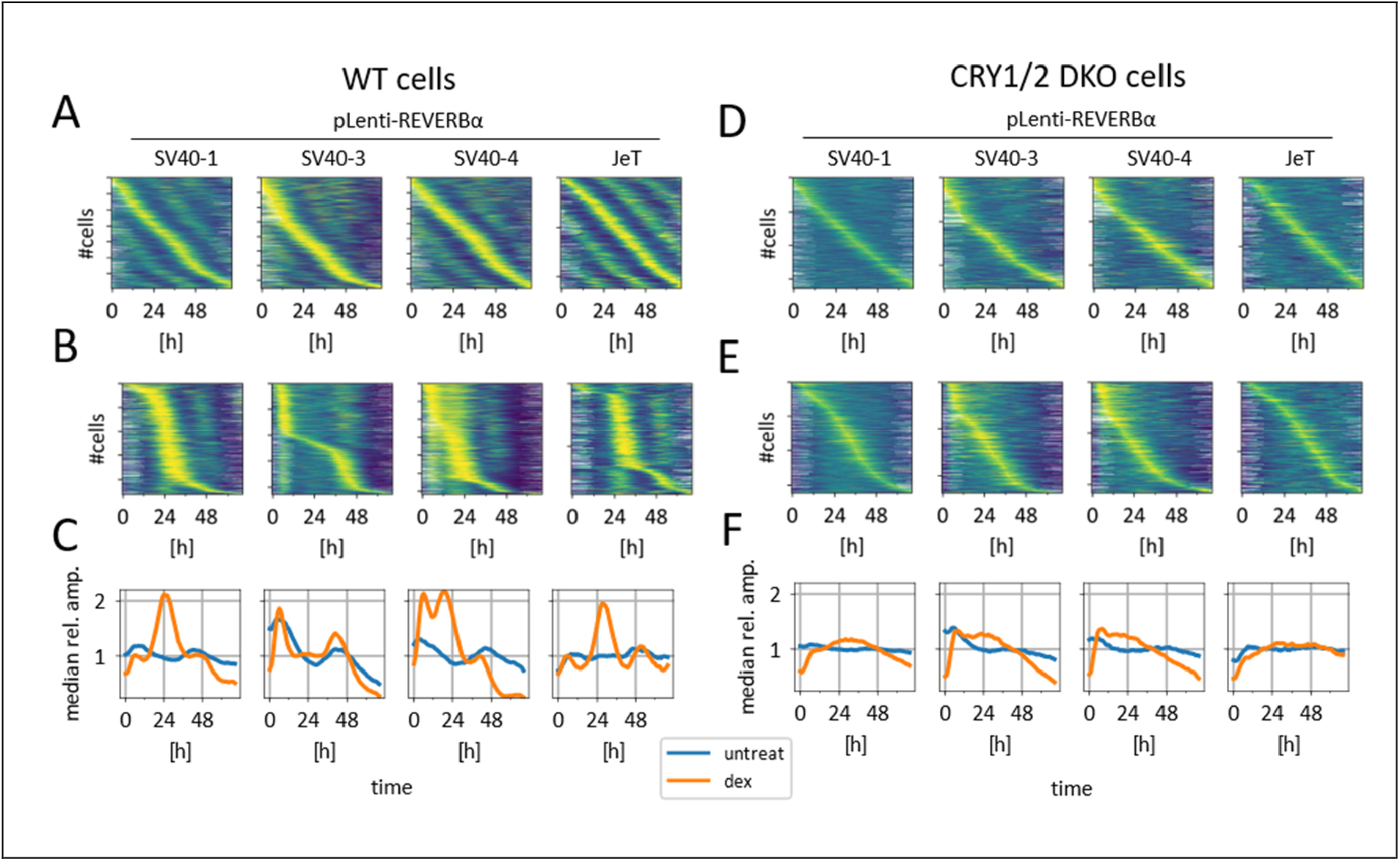
Reporter oscillations depend on a functional circadian clock. **(A, B)** Heat maps showing normalized fluorescent signal oscillation from the indicated pLenti-REVERBα circadian reporters in individual wild-type cells, either unsynchronized (A) or treated with dexamethasone (B). **(C)** Population median signal for all time-series shown in (A) and (B). **(D, E)** Heat maps of CRY1/CRY2 double knock-out single cells, either unsynchronized (D) or dexamethasone-treated (E). **(F)** Population median signal of all time-series shown in (D) and (E). Timeseries were normalized to their median signal, filtered for the 50% most rhythmic time series of each population. Traces are sorted by the timepoint of maximal signal. Y-axis ticks indicate every 250th cells.

### Determination of circadian parameters in single cells

To evaluate the ability of the lentiviral reporters to accurately report circadian rhythms in individual cells, we compared their expression dynamics with those of the full-length reporter and with the circadian proteins CRY2 and PER1 within the same cells. To this end, we replaced the Venus protein in the lentiviral reporters with the red fluorescent protein mScarlet-I3, and transduced them into (1) a clone that expresses the full-length reporter coupled to Venus, and (2) knock-in cells that express yellow fluorescent PER1-mClover3 or CRY2-mClover3 fusion proteins from the endogenous locus (generated as described (Gabriel et al. 2024), see Method section for details). We recorded fluorescence in both channels over the course of three days and extracted nuclear fluorescence time series for each cell (examples shown in Fig. 3A-F). Sorting the time-series by peak expression time again revealed oscillatory patterns (Fig. 3G), that were remarkably similar to those of the full-length REVERBα reporter (Fig. 3H), and the knock-in fusion proteins (Fig. 3I). We analyzed single-cell time series using wavelet analysis (Schmal et al. 2022), extracting circadian period, amplitude, average wavelet power (which serves as a measure of rhythmicity), and phase at 36 hours after the start of the recording. We then compared phase, amplitude and period of the most rhythmic time series (average wavelet power > 7 for both reporter types).

**Figure 3:**
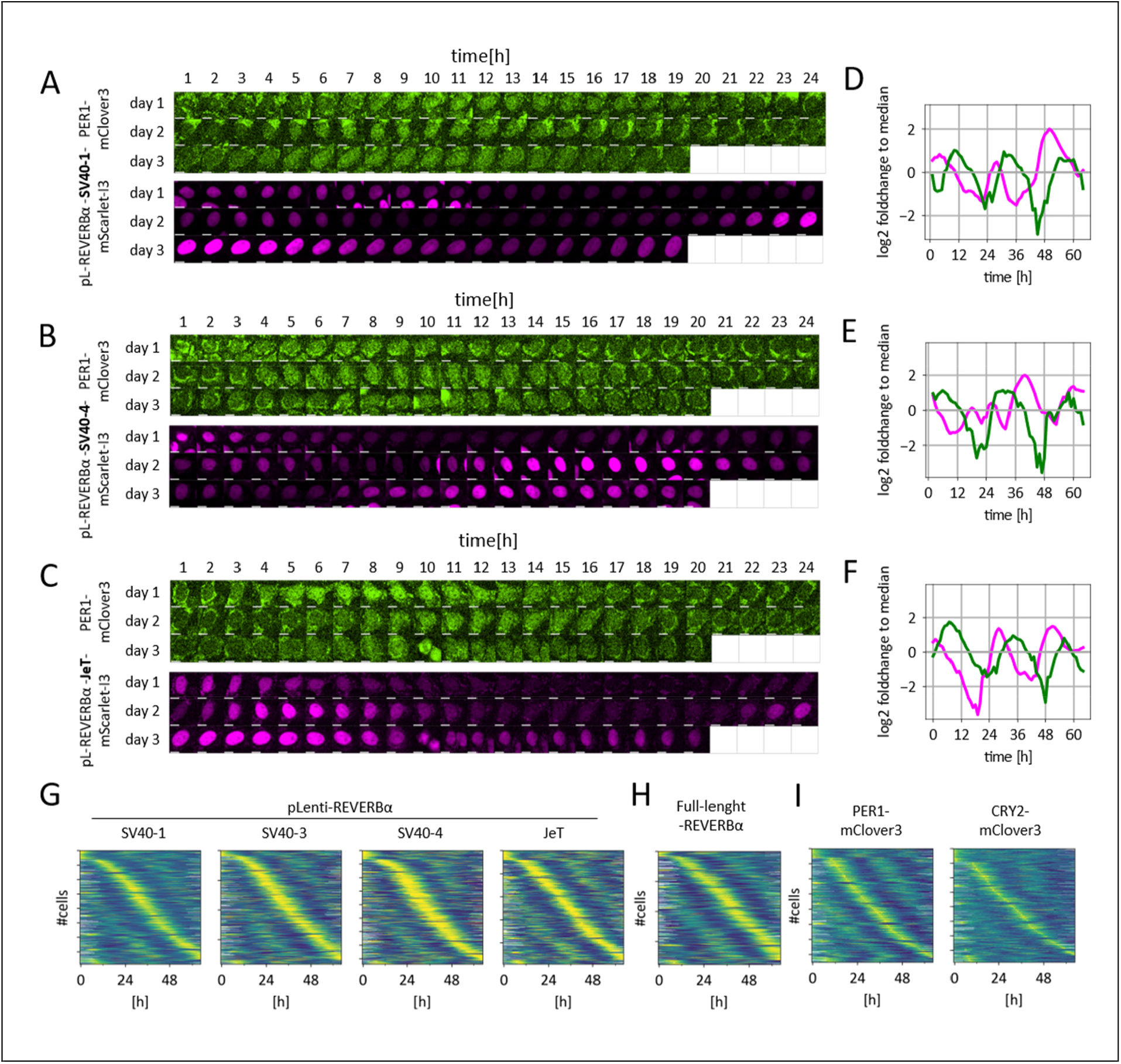
Generation of cells with two circadian fluorescence reporters. **(A, B, C)** Dual-color time-lapse recordings of nuclear fluorescence in single cells (background subtracted). Green: PER1-mClover3 fusion protein. Magenta: pLenti-REVERBα-mScarlet-I3 with SV40-1 (A), SV40-4 (B) or JeT (C) promoter. Scalebar: 10 µm. **(D, E, F)** Mean nuclear fluorescence signals plotted as log2 fold change relative to the median of each time series. **(G, H, I)** Heat maps of normalized fluorescent signal oscillations from the indicated pLenti-REVERBα circadian reporters (G), the full-length REVERBα-Venus reporter (H), and CRY2- or PER1-mClover3 fusion proteins in unsynchronized dual-reporter single cells (I). All time series were filtered to retain the 50% most rhythmic time series within each group and sorted by time of maximal signal. Y-axis ticks indicate every 250th cells.

While the circadian periods measured using SV40 promoter-containing reporters correlated significantly with those derived from the full-length construct (Spearman correlation coefficient: 0.35-0.53), correlations with PER1- or CRY2-mClover periods were weaker (correlation coefficients 0.05 – 0.29) (Fig. 4A). Conversely, the JeT promoter-containing construct showed stronger correlation with PER1 than with the full-length REVERBα reporter (Fig. 4A).

**Figure 4:**
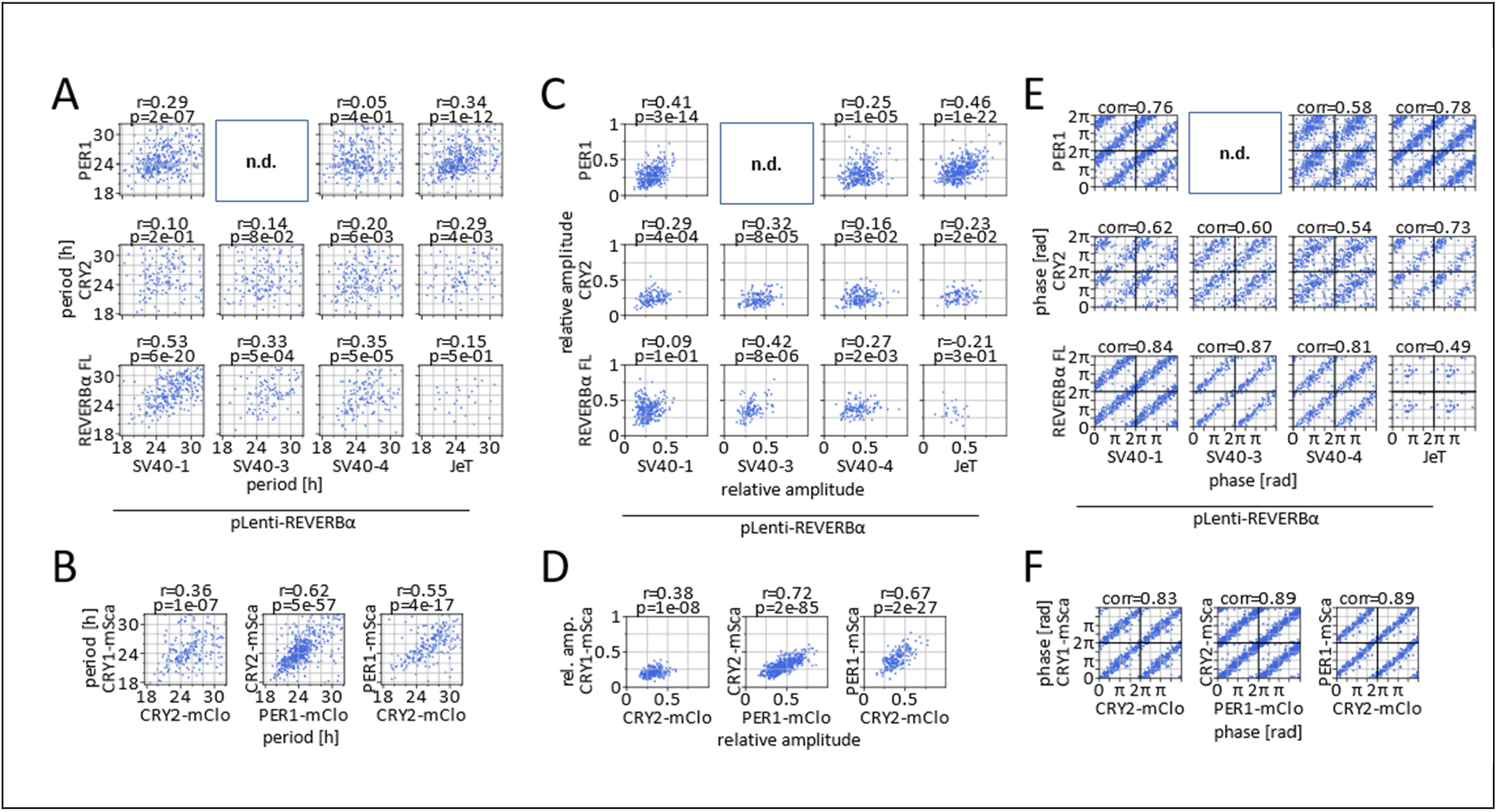
Correlation of circadian dynamics parameters between different reporters in single cells. **(A, B)** Correlation of circadian periods, **(C, D)** relative amplitudes, and **(E, F)** circadian phases (36 h after recording onset), calculated from time series of the two indicated circadian fluorescent reporters expressed in the same single cells. Only cells exhibiting rhythmic signals in both reporters were included in the analysis. Phases are double-plotted along both axes. p, r: Spearman correlation, corr: circular moment of phase differences, FL: full-length REVERBα–Venus reporter.

In general, the relative amplitudes of the lentiviral reporters align more closely with those of the fluorescence-tagged repressors than with the full-length reporters. Notably, the similarity in period and amplitude between the JeT-containing reporter and the PER1 protein reporter resembled that observed between CRY1- and CRY2 protein in double knock-in cells, though it was less than that seen between PER1 and CRY2 in double knock-in cells (Fig. 4B and D). Phases derived from the SV40 promoter-containing reporters more closely matched those of the full-length reporter, whereas the JeT-driven variant showed greater similarity with the endogenous fusion proteins (Fig. 4E). In particular, the JeT and SV40-1 variants showed notable phase correlations with the endogenous proteins (Fig. 4E-F).

Taken together, the lentiviral REVERBα reporters are suitable to accurately determine circadian dynamics parameters such as circadian phase and, to a lesser extent, circadian period and amplitude in individual cells. Among the constructs tested, the lower-expressing JeT-driven reporter outperformed those driven by SV40 promoters.

## Discussion

Here, we report on the creation of a lentiviral fluorescent reporter of circadian rhythms, based on the murine *Nr1d1/REVERBα* gene. While retaining primarily BMAL1 bound sites was sufficient to allow for regulation by the circadian clock machinery, the expression levels were significantly lower than those of the full-length reporter that includes a much larger proportion of the genomic locus. This suggests that CLOCK/BMAL1 binding alone does not drive robust transcription of the target genes, but rather modulates their expression dynamics through timed inhibition and enhancement of transcription, (e.g. via chromatin remodeling (Trott and Menet 2018)), thereby aligning gene expression with circadian rhythms.

Adding constitutive promoters enhanced the reporter signal without overriding it. As shown in Fig. 2, oscillations are clearly present in the wild-type cells but not in knock-out cells, demonstrating that these oscillations are primarily governed by the circadian clock rather than the constitutive promoters. Nevertheless, the constitutive promoters can still influence signal dynamics; for example, acute dexamethasone-induced increases in signal were observed for the variants SV40-3 and SV40-4 (Fig. 3). Therefore, adequate controls are necessary to rule out artifacts introduced by these promoters.

Surprisingly, the correlation of periods derived from different reporters was relatively low, even in double knock-in cells expressing two tagged clock proteins. However, it is unlikely that e.g. CRY1 and CRY2 oscillate with significantly different periods in the same cell. Rather, day-to-day period variability and/or transcriptional noise likely introduce uncertainty in period calculation, producing the apparent divergence. When analyzing comparatively short time series (2-3 days) differences in one or two measurement values may greatly affect the calculated period. Period estimates between reporters tend to correlate more strongly in longer recordings.

In contrast, the JeT and SV40-1 containing reporters proved particularly effective in estimating circadian phases at the single-cell level. This is demonstrated by the high correlation with phases derived directly from circadian core clock proteins (CRY2 and PER1), with the level of agreement almost reaching that of two core clock proteins (Fig. 4). Accurate phase determination in individual cells allows researchers to determine whether a cell’s response to a stimulus depends on its circadian phase. This could help identify, for example, the most efficient times for chemotherapeutics (Ector et al. 2024), or optimal windows for cell differentiation (Zhang et al. 2022).

The ease with which the lentiviral REVERBα reporters can be delivered should enable visualization of single-cell circadian rhythms in a broad range of cell types using a simple protocol, as clonal selection is not required. This also extends to primary cells, thus enabling screening for circadian phase-dependent drug or toxin effects in complex, physiologically relevant cell culture systems. We hope that our lentiviral circadian reporters will open up the field of single-cell circadian rhythms and their significance to a wide range of research questions and applications.

## Methods

### Cells

U-2 OS (RRID:CVCL_0042, human, female, ATCC HTB-96) cells were cultured in DMEM supplemented with 10% FBS (Life, lot 2453915), 25 mM HEPES and penicillin/streptomycin at 37°C and 5% CO_2_. The U-2 OS CRY1/2 double knock-out cells were described before (Bording et al. 2019). U-2 OS cells stably expressing an NR1D1::VNP fusion protein were provided by Michael Brunner (University of Heidelberg). A single subclone was used for experiments. To allow automated detection of nuclei, all clones were transduced to express a histone-2B-iRFP720 fusion protein, which results in nuclear expression of the infrared protein miRFP720 (RRID:Addgene_128961), and cells were sorted for high expression by FACS.

### Generation of knock-in cells

The generation and validation of single and double circadian reporter U-2 OS knock-in cells using CRISPR has been described in detail in (Gabriel et al. 2021; Gabriel et al. 2024). In brief, cells are transfected by electroporation with three plasmids containing (1) Cas9 and sgRNA targeting CRY2 or PER1 downstream of the stop codon (RRID:Addgene_189989, RRID:Addgene_189987), (2) a template for homology directed repair (HDR) that will result in a fluorophore (mClover3) integrated C-terminally (RRID:Addgene_189979, RRID:Addgene_189983), and (3) an inhibitor of non-homologous end-joining to enhance HDR (pCAG-i53bp, a gift from Ralf Kuhn and derived from RRID:Addgene_74939). After positive selection, negative selection and CRE-mediated removal of selection cassettes, CFP- and CD4-negative sorted cells were clonally expanded and screened for fluorescent clones. Knock-in specificity is controlled by PCR and by using shRNA targeting either PER1 or CRY2, and intactness of the circadian clock is tested by analysis of signals from a transduced mBmal1-promoter driven luciferase reporter (Maier et al. 2009). For this study, CRY2-mClover3 and PER1-mClover3 single knock-in cells were generated from WT U-2 OS cells, CRY1-mScarlet-I/CRY2-mClover3 cells from CRY1-mScarlet-I single knock-in cells (described in (Gabriel et al. 2021)), PER1-mScarlet-I/CRY2-mClover3 from PER1-mScarlet-I clone #1, and CRY2-mScarlet-I/PER1-mClover3 from CRY2-mScarlet-I clone #1 single knock-in cells (both described in PNAS).

### Construction of the lentiviral REVERBα reporters

The basic version of pLenti6-puromycin-mREVERBα-Venus-NLS-Pest (pL6-RAV-ori) was synthesized by VectorBuilder. The reporter sequence was based on the mREVERBα-Venus reporter plasmid published by (Nagoshi et al. 2004) and the murine *Nr1d1* gene (NC_000077.7). It included the following parts: 633 bp upstream of the transcription start site (TSS), exon-1, intron-1(bp 1-22, 528-971, 1480-2051), exon-2(bp -14 - 138) coupled to Venus - SV40-Nuclear localization signal (NLS) with destabilization domain from Ornithin decarboxylase, and the 3’UTR (last 288 bp of exon 8).

The JeT promoter sequence (Tornoe et al. 2002) was synthesized (MWG Eurofins) with flanking restriction sites for KpnI and EcoRI. The SV40 promoter or minimal promoter sequence was amplified from a lentiviral expression plasmid using primers creating overhangs for restriction enzymes. The promoters were inserted either into the EcoRV site ∼ 500 bp upstream of the TSS or the KpnI site ∼70 bp downstream of the TSS by restriction enzyme cloning and sequences verified by Sanger sequencing. Promoter sequences are listed in Supplementary Table S1.

The fluorophores mScarlet-I3, mClover3, SYFP2, miRFP713 Electra2 were amplified from Addgene plasmids #197230, #189763, #179479 and #179441 and used to replace the Venus protein by restriction enzyme cloning using XbaI and MluI. All plasmid and the corresponding sequences can be obtained from Addgene. (RRID:Addgene_240105-240118).

### Lentivirus production

HEK293-T cells were transiently transfected in a T75 flask with 8.6 μg lentiviral expression plasmid, 6 μg psPAX2, and 3.6 μg pMD2G (gift from Trono lab, RRID:Addgene_12259 and RRID:Addgene_12260) packaging plasmid using the CalPhos Mammalian Transfection Kit (Takara). The next day, culture medium was replaced with 12.5 mL of medium (DMEM for U-2 OS cell, M199 without ECGF for HUVECs), and lentiviral supernatant was collected after 24 and 48 h. The combined supernatant was passed through a 0.45-μm filter (Filtropur S 0.45) and either used directly or stored in aliquots at -80°C. For transduction, cells were seeded into lentivirus-containing supernatant supplemented with 8 μg/mL protamine sulfate. The next day, lentivirus-containing supernatant was aspirated and cells were cultured in complete culture medium for another 24 hours before antibiotic selection of transduced cells.

### Live-cell fluorescence microscopy

For microscopy, cells were seeded on glass bottom #1.5H-N 96-well plates (Cellvis, USA) coated with 50 µg/ml human serum fibronectin (Merck, Germany). Cells were left untreated or synchronized with 100 nM dexamethasone 3 h prior to recording for 20 min, washed with and supplemented with fresh medium. Image acquisition was done in Flurobrite medium (GIBCO) supplemented with 2% FBS, 1:100 PenStrep, and 1x GlutaMax at 37 °C and 5% CO2. Imaging was performed on a Nikon Widefield Ti2 equipped with a pco.edge 4.2 USB sCMOS camera (Excelitas pco, Germany) and an incubator with environmental control (OKO Labs bold line, Italy). Illumination was provided by a SpectraX LED system (Lumencor, USA) on the following microscope-presets:The following light sources (LEDs) and emission filters and settings were usually used for the different channels: GFP (mClover3, Venus), excitation 475/28 nm, 57 mW, 50 % intensity, 200ms (Fig. 1D, 3, 4A, 4C), emission 519/26 nm; YFP (mClover3, Venus): excitation 511/16 nm, 12.3 mW, 25% intensity, 100 ms, (Fig. 2) / 50% intensity, 800 ms (Fig. 1C), 30% intensity, 1s (Fig. 4B, 4D), emission 540/30 nm; RFP (mScarlet-I, mScarlet-I3) excitation 555/28 nm, 145 mW, 40% intensity (Fig.2) / 50% intensity, 100 ms (Fig. 1D, 3, 4A, 4C) / 50% intensity, 800 ms (Fig. 4A, 4C: JeT-REVERBα-FL) / 12% intensity, 2s (Fig. 4B,4D), emission 642/80 nm; iRFP: excitation: 625/22 nm, 38.9 mW, 100% intensity, 200ms, emission 697/50 nm. The used dichroic mirrors for presets “GFP”, “RFP”, “iRFP” were a quad dichroic mirror ET435/33, 526/20, 595/38, 695/63 and for “YFP” a quad dichroic mirror ET475/25, 537/30, 644/92, 806/100. Objectives: Objectives: 20x Plan Apo, NA 0.75, WD 1000 µm (Nikon, Japan) (Fig. 2), 40x Plan Apo, NA 0.95, WD 250 μm (Nikon, Japan) (Fig. 1,3,4). Time-lapse images were acquired with a 1h interval for up to three days days.

### Single cell tracking

Cell tracking was performed automatically by the help of ImageJ-Fiji (Schneider et al. 2012) and the TrackMate plugin. Two macros were used for preprocessing and for segmentation/tracking, respectively.

Preprocessing: For uneven illumination correction, 100 images from buffer only wells were used to generate relative illumination patterns for each channel, and every image was corrected for non-uniform illumination by dividing pixel by pixel by the previously generated patterns. The background of the illumination-corrected RFP and YFP images was determined as the modal value of a gaussian filter (size=50) smoothened image and subtracted.

Segmentation/Tracking: The trackmate plugin of Fiji was used for segmentation and tracking of nuclei (Ershov et al. 2022). The iRFP channel was used for segmentation of nuclei in each image using the StarDist module (Schmidt et al. 2018), and objects were filtered by size (min = 3 µm). Subsequently, objects were tracked using the LAP algorithm (maximal range = 30 µm, no gaps allowed). Finally, the mean fluorescence intensity for each tracked nucleus for each time point was extracted from all channels. After cell division, tracking continued with one daughter cell, while the other daughter cell was considered a newly emerging object.

For quality control of tracking, we used a Python Script as described in (Gabriel et al. 2024) and available from GitHub that sorts out most mis-segmented cells and tracking errors. Fluorescence intensities at cell division and subsequent time points were linearly extrapolated from neighboring time points, because detachment of dividing cells produced fluorescence artifacts. Visual inspection of a set of 50 tracks confirmed correct segmentation and tracking for 48 of these tracks.

### Data analysis

RFP-channel fluorescence values were compensated for Venus bleed-through into the RFP channel in REVERBα-FL expressing cells (Fig. 3, 4). Signals whose median fluorescence signal did not surpass that of 99% of non-fluorescent cells were filtered out.

We used the python based pyBOAT toolkit (Schmal et al. 2022) to analyze rhythmicity in single cell time series by wavelet analysis. We allowed periods to fall between 18-32 h, detrended raw data using sinc_detrend (T_c=50), calculated ridge_data with power_threshold=0 and smoothing_wsize=4, and extracted instantaneous phases at indicated timepoints. Ridge data was cropped by disregarding values from the first and last 9 present timepoints before calculating average wavelet power as the mean of the maximal power of the ridge, (average) period as the median of instantaneous period, and (average) absolute amplitude as the mean of instantaneous amplitudes from the remaining timepoints. Relative amplitudes were calculated by dividing absolute amplitudes by mean expression intensity. We defined time-series as rhythmic if average wavelet power of a 3-days timeseries surpasses 7, based on analysis of negative control datasets (autofluorescence and tracking marker intensity time series). Additionally, we excluded times series that were considered rhythmic, but whose calculated period exactly matched the input limits (18-32 h), as these were most likely to contain oscillations longer or shorter than these limits. For data from dexamethasone treated cells, the first 18 hours of recording were omitted for wavelet analysis.

Correlation analysis were done for data from those cells which had rhythmic signals from both reporters. For correlation of phase data, we calculated phase differences of the two reporters for each cells at a single time-point and calculated the circular moment of the phase differences using the *circmoment* method of the python module astropy.stats.circstats (Price-Whelan et al. 2018). This tests whether the two reporters have a defined phase relation, considering that they can be phase-shifted. Periods correlation was tested by Spearman correlation. All plots were generated using the *pandas, scikit-image* and *matplotlib* python libraries.

### AI

Artificial intelligence was used to enhance language and readability of the text. The text was checked by the authors for correctness afterwards.

## Data and material availability

Fluorescence time series data to reproduce the figures are provided in supplementary tables S2-S3. Imaging raw data is available from the authors upon request. All plasmids are available from Addgene (Plasmids 240105-240118). All code is available at https://github.com/Kramer-Lab/Gabriel-et-al_2025_CircLentiReporter)

## Acknowledgement

We thank Michael Brunner and Emi Nagoshi for providing material. We acknowledge the support of the FCCF at the German Rheumatism Research Center. We also thank the Advanced Medical Bioimaging Core Facility (AMBIO) of the Charité for assistance with the acquisition of imaging data. This work was funded by the German Research Foundation (DFG) - project number 278001972 - TRR 186.

## Author contributions

C.H.G. and A.K. conceived the project. C.H.G. designed and performed experiments with assistance from L.L. and J.A. C.H.G. developed image analysis pipelines and performed quantitative data analysis. A.K. supervised the study and secured funding. C.H.G. wrote the manuscript with input from all authors. All authors reviewed and approved the final version of the manuscript.

## Supplementary Material

**Supplementary Videos SV1: Time-lapse microscopy of U-2 OS cell transduced with indicated lentiviral REVERBα-mScarlet-I3 reporter**. Scale bar=20 µm.

**Supplementary Table S1:** Sequences of the constitutive promoters.

|  |  |  |  |
| --- | --- | --- | --- |
| JET Promoter | GAATTCGGGC | GGAGTTAGGG | CGGAGCCAAT |
|  | CAGCGTGCGC | CGTTCCGAAA | GTTGCCTTTT |
|  | ATGGCTGGGC | GGAGAATGGG | CGGTGAACGC |
|  | CGATGATTAT | ATAAGGACGC | GCCGGGTGTG |
|  | GCACAGCTAG | TTCCGTCGCA | GCCGGGATTT |
|  | GGGTCGCGGT | TCTTGTTTGT | GGATCCCTGT |
|  | GATCGTCACT | TGACA |  |
| SV40<br>Enhancer/Promoter | GTGTGTCAGT | TAGGGTGTGG | AAAGTCCCCA |
|  | GGCTCCCCAG | CAGGCAGAAG | TATGCAAAGC |
|  | ATGCATCTCA | ATTAGTCAGC | AACCAGGTGT |
|  | GGAAAGTCCC | CAGGCTCCCC | AGCAGGCAGA |
|  | AGTATGCAAA | GCATGCATCT | CAATTAGTCA |
|  | GCAACCATAG | TCCCGCCCCT | AACTCCGCCC |
|  | ATCCCGCCCC | TAACTCCGCC | CAGTTCCGCC |
|  | CATTCTCCGC | CCCATGGCTG | ACTAATTTTT |
|  | TTTATTTATG | CAGAGGCCGA | GGCCGCCTCT |
|  | GCCTCTGAGC | TATTCCAGAA | GTAGTGAGGA |
|  | GGCTTTTTTG | GAGGCCTAGG | CTTTTGCAAA |
| SV40 Enhancer | GTGTGTCAGT | TAGGGTGTGG | AAAGTCCCCA |
|  | GGCTCCCCAG | CAGGCAGAAG | TATGCAAAGC |
|  | ATGCATCTCA | ATTAGTCAGC | AACCAGGTGT |
|  | GGAAAGTCCC | CAGGCTCCCC | AGCAGGCAGA |
|  | AGTATGCAAA | GCATGCATCT | CAATTAGTCA |
|  | GCAACCATAG | TCCCGCC |  |

**Supplementary Table S2**: Source data to Figures 1, 3, 4 (separate file)

**Supplementary Table S3**: Source data to Figure 2 (separate file)

